# Chromatin Buffers Torsional Stress During Transcription

**DOI:** 10.1101/2024.10.15.618270

**Authors:** Jin Qian, Lucyna Lubkowska, Shuming Zhang, Chuang Tan, Yifeng Hong, Robert M. Fulbright, James T. Inman, Taryn M. Kay, Joshua Jeong, Deanna Gotte, James M. Berger, Mikhail Kashlev, Michelle D. Wang

## Abstract

Transcription through chromatin under torsion represents a fundamental problem in biology. Pol II must overcome nucleosome obstacles and, because of the DNA helical structure, must also rotate relative to the DNA, generating torsional stress. However, there is a limited understanding of how Pol II transcribes through nucleosomes while supercoiling DNA. In this work, we developed methods to visualize Pol II rotation of DNA during transcription and determine how torsion slows down the transcription rate. We found that Pol II stalls at ± 9 pN·nm torque, nearly sufficient to melt DNA. The stalling is due to extensive backtracking, and the presence of TFIIS increases the stall torque to + 13 pN·nm, making Pol II a powerful rotary motor. This increased torsional capacity greatly enhances Pol II’s ability to transcribe through a nucleosome. Intriguingly, when Pol II encounters a nucleosome, nucleosome passage becomes more efficient on a chromatin substrate than on a single-nucleosome substrate, demonstrating that chromatin efficiently buffers torsional stress via its torsional mechanical properties. Furthermore, topoisomerase II relaxation of torsional stress significantly enhances transcription, allowing Pol II to elongate through multiple nucleosomes. Our results demonstrate that chromatin greatly reduces torsional stress on transcription, revealing a novel role of chromatin beyond the more conventional view of it being just a roadblock to transcription.

---

During transcription elongation, RNA polymerase (RNAP) and DNA must rotate relative to each other as RNAP translocates along the DNA. Because the nascent RNA is often associated with large protein complexes (such as ribosomes in prokaryotes and spliceosomes in eukaryotes)^1^, RNAP cannot rotate freely, and thus, the DNA must rotate. This obligatory relative rotation, dictated by the helical nature of DNA^2, 3^, imposes one turn for every 10.5 nt transcribed. Consequently, RNAP tightly couples its translocation and rotation, making it concurrently a linear motor to generate force to translocate and a rotary motor to generate torque (torsion) to rotate^4, 5^. These inherent motor properties are crucial for gene expression, enabling RNAP to use mechanical energy to overcome roadblocks and reconfigure DNA structures and topology^6, 7^. Notably, the rotational motion leads to DNA supercoiling and torsion in the DNA, which, in turn, regulates transcription^6-13^.

Recent studies have highlighted the diverse critical roles of transcription-generated DNA supercoiling in cellular functions^14-18^. DNA supercoiling is present across the genome and regulates gene expression, even in the presence of native topoisomerases^19-24^. DNA supercoiling generated from the transcription of one gene can also regulate the expression of other genes in the vicinity^7, 9, 25, 26^. Moreover, DNA supercoiling accumulation during a head-on conflict of transcription and replication has been suggested to promote persistent replication stress and subsequent DNA damage^27-29^. It is worth noting that because torsion in DNA can act over a distance, this consequence may occur well before any physical encounter of RNAP with the replisome^27, 30^. Furthermore, transcription-generated supercoiling has also been found to significantly change DNA topology by regulating 3D genome knotting, folding, and loop formation^31^.

Torsion (torque) in the DNA originates from RNAP’s ability to rotate the DNA while working against the resistance of the DNA to rotation. Yet our understanding of how RNAP generates torque and works against torsion is limited. Previously, we found that *E. coli* RNAP is a powerful rotary motor capable of generating sufficient torque to melt DNA^10, 11^. Whether the eukaryotic RNAP Pol II can do the same remains unknown.

Importantly, Pol II must transcribe through nucleosomes; thus, transcription through chromatin under torsion represents a fundamental problem in biology. Despite the significance of this problem, little is known about how Pol II transcribes through chromatin while supercoiling DNA. Chromatin is commonly perceived as a roadblock to transcription elongation, but this perception must be revisited, especially in the context of transcription under torsion. However, the complexity of this problem poses significant challenges to experimental investigation and requires new methodologies and conceptualizations to gain mechanistic understanding. In this work, we have enabled the investigation of transcription through chromatin under torsion and provided unique and quantitative insights into this process.

## Visualizing Pol II rotation of DNA

The fundamental cause of transcription-generated torsion is RNAP rotation of DNA. Understanding this rotation requires a real-time method for visualizing this rotational motion when transcription occurs at a specified torsional stress level. Previously, the rotational motion for *E. coli* RNAP was detected/inferred via the rotation of a bead^32^, the rotation of a DNA origami attached to the DNA^33^, or RNAP-generated DNA supercoiling^10, 11^. However, these methods either do not allow on-demand control of the torsional stress or they require signal conversion to track the rotation.

Here, we present an approach to directly track Pol II-generated rotation of DNA at high spatial and temporal resolution using an angular optical trap (AOT). A defining feature of the AOT is its trapping particle, a nanofabricated birefringent quartz cylinder (Fig. 1a; Supplementary Fig. 1), which serves as a handle for simultaneous measurements of force, extension, torque, and rotation of DNA^34-37^. The angular orientation of the quartz cylinder can be detected at an exceedingly high angular and temporal resolution^38, 39^ and thus can be used to visualize how Pol II induces DNA rotation.

**Figure 1.**
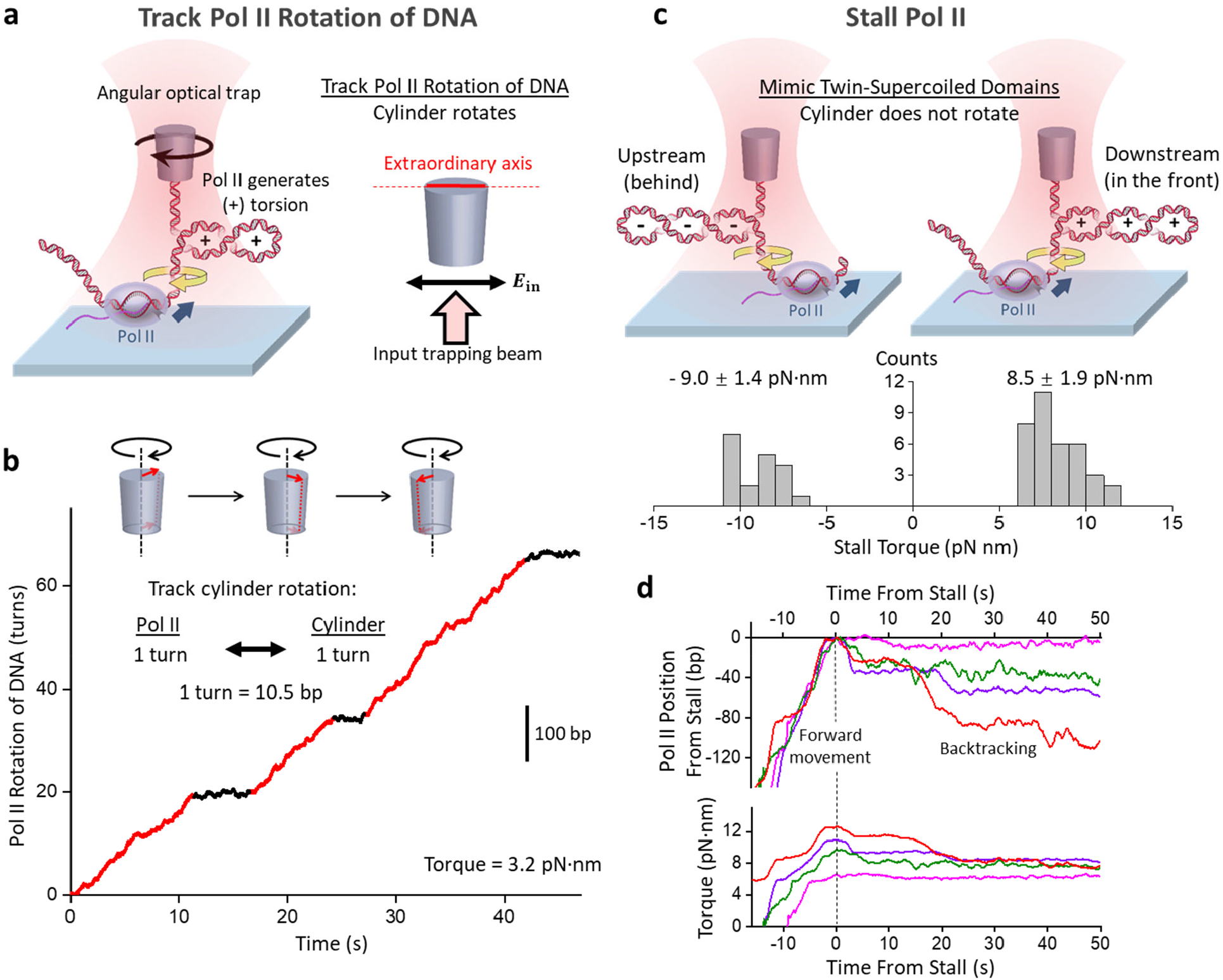
Visualizing Pol II rotation and stalling under torsion. **a**. An experimental configuration to track Pol II rotation of DNA during transcription using an angular optical trap (AOT). Pol II is torsionally anchored on the surface of a coverslip, and its downstream DNA is torsionally attached to the bottom of a quartz cylinder, which is trapped by the AOT. Pol II elongation generates (+) supercoiling in the downstream DNA. The cylinder rotates to follow Pol II rotation to maintain a constant torque in the DNA. The right panel shows that the extraordinary axis of the cylinder tends to align with the linear polarization of the input trapping beam. The orientation of this axis is accurately detected and informs the angular orientation of the DNA attached to the cylinder. **b**. A representative real-time trajectory of Pol II rotation of DNA during transcription under a +3.2 pN·nm resistance torque. The scale bar shows the conversion to base pair position of Pol II (or nucleotides transcribed). Regions of Pol II steady rotation are marked red, and regions of pausing are marked black. See Supplementary Movie 1 for the corresponding video. **c**. Pol II stall torque measurements using the AOT. The two cartoons illustrate the experimental configurations to stall Pol II against (-) torsion upstream (left) and (+) torsion downstream (right), mimicking the “twin-supercoiled-domain” model of transcription. The measured stall torque distributions are shown beneath the corresponding cartoon, with the mean and SEM indicated. **d**. Representative traces of Pol II backtracking upon stalling under (+) torsion. Different colors represent different traces. Pol II forward translocates and stalls under increased (+) torque. Upon stalling, Pol II backtracks, as evidenced by the reverse motion.

To apply the AOT to Pol II, we torsionally constrained Pol II to the coverslip surface of a sample chamber and its downstream DNA to the bottom of the cylinder (Fig. 1a; Supplementary Figs. 2 & 3). Pol II translocation leads to positive (+) DNA supercoiling and buckles the DNA to form a plectoneme (Supplementary Fig. 4). Once the torque reaches a desired value, we rotate the cylinder to follow Pol II rotation of DNA, thus limiting further torsion buildup and thereby maintaining a constant torque. At this point, for each turn of Pol II rotation of the DNA (10.5 bp transcribed), the cylinder also rotates by a turn (Methods). Because the cylinder’s extraordinary axis angle can be tracked at exceedingly high spatial and temporal resolution, this method provides unprecedented resolution of the Pol II rotational motion.

**Figure 2.**
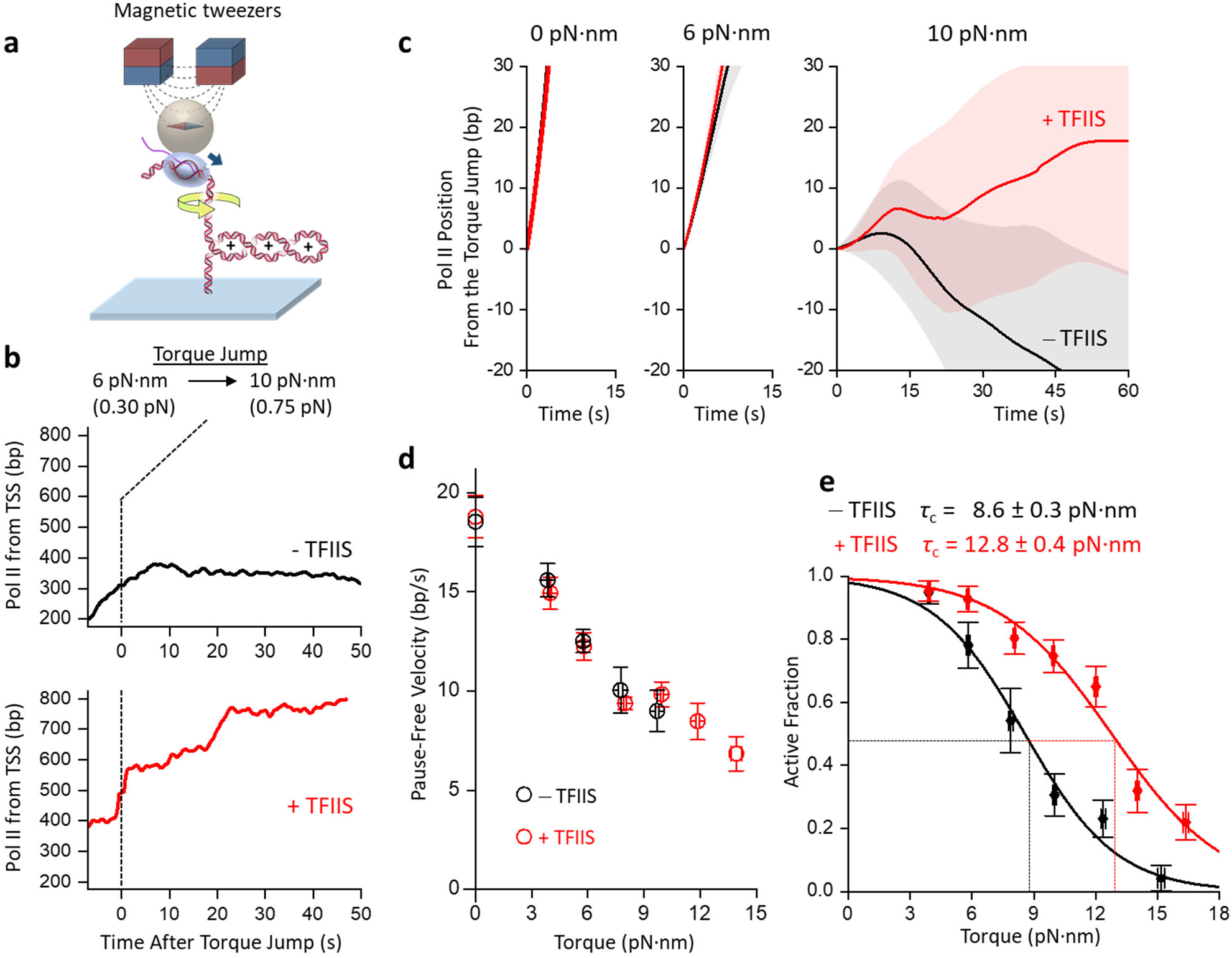
TFIIS up-regulates Pol II’s ability to transcribe under torsion. **a**. Experimental configuration to monitor transcription activity under torsion via magnetic tweezers. Pol II is torsionally anchored to the surface of a magnetic bead, while its downstream DNA is torsionally anchored to the surface of a coverslip. The magnetic bead can be used to manipulate the torsional state of the DNA and the force on the DNA. **b**. Example traces of Pol II elongation during a torque-jump experiment. Pol II transcription starts at a torque of low value before jumping to a higher value. Since the DNA is buckled, the force on the DNA informs the torque in the DNA (Supplementary Fig. 4d). The two plots show the Pol II position from the transcription start site (TSS) without and with TFIIS. **c**. The mean trajectory of Pol II elongation under different torques. All traces were aligned at the start of the torque jump (*t* = 0), with the shaded region for each curve representing 30% of the standard deviation. Without TFIIS: *N* = 87, 72, and 42 for 0, 6, and 10 pN·nm, respectively. With TFIIS: *N* = 36, 72, and 61 for 0, 6, and 10 pN·nm, respectively. **d**. Torque-velocity relation. Pause-free velocities are shown without and with TFIIS. The error bars represent the SEM, with each data point collected from *N* = 27-87 traces. **e**. Active fraction after the torque jump. The active fraction is the fraction of traces remaining active at a given torque. Each curve is fit with a decaying function (Methods) to obtain the critical torque *τ*_c_, at which 50% of traces are active. The fit values and uncertainties without and with TFIIS are also shown.

The example trace in Fig. 1b shows Pol II rotation of DNA under a relatively low 3.2 pN·nm resisting torque, mimicking an *in vivo* condition when topoisomerases can almost keep up with transcription. As shown, as Pol II progresses along the DNA, it exhibits steady rotational motion (∼ 2 turns/s), interspersed by pauses. This rotational motion is visualized in Supplementary Movie 1. These Pol II rotational behaviors are reminiscent of the translocational motions of RNAP from previous studies^40-46^, reflecting the translocation-rotation coupling of RNAP. The direct visualization of the Pol II rotational motion highlights the inherent nature of rotational motion during transcription and invites the investigation of the inevitable consequences of rotational motions.

### Pol II is a powerful rotary motor

*In vivo*, Pol II rotation is thought to be restricted due to the viscous drag of large molecular machinery, such as spliceosomes, associated with the nascent RNA^47^. Consequently, transcription accumulates positive (+) supercoiling in front of Pol II and negative (-) supercoiling behind when supercoiling cannot be fully dissipated at the ends of the DNA, leading to the well-known “twin-supercoiled-domain” model of transcription^48^. Both the (+) supercoiling in front of Pol II and the (-) supercoiling behind it create resistance to transcription. These torsional buildups can occur even in the presence of topoisomerases^19-23^, suggesting that topoisomerases cannot always keep up with transcription. Especially when there is a transient shortage of topoisomerases near an active gene, torsion in the DNA can build up, leading to Pol II stalling. The torque at which Pol II stalls determines Pol II’s torsional capacity to work against the torsional stress and modulate the DNA topology. However, it is unknown how much torque Pol II can generate before stalling.

To measure Pol II’s stall torque, we adapted a method we previously used for stall torque measurements of *E. coil* RNAP using an angular optical trap (AOT)^10, 11^. Instead of rotating the cylinder as in Fig. 1a, we restricted the cylinder rotation, allowing Pol II to build up torsion and eventually stall (Fig. 1c). During this process, we monitored how Pol II moves along the DNA and generates torsion in real-time (Fig. 1d). We investigated two stalling configurations: Pol II accumulation of (-) supercoiling behind or (+) supercoiling in the front and found that the stall torque magnitudes are similar in these two configurations despite being opposite in rotational sense: -9.0 ± 1.4 pN·nm (mean ± SD) and 8.5 ± 1.9 pN·nm, respectively. For comparison, the DNA melting torque is around -10 pN·nm^36, 49^, suggesting that Pol II is a torsional motor almost sufficiently powerful to melt DNA.

### TFIIS enhances Pol II’s torsional capacity

Upon stalling under torsion, we observed that Pol II often reverses its movement and moves backward by as much as 100 bp or more while relaxing the torsional stress (Fig. 1d). This reverse motion is consistent with Pol II backtracking, during which Pol II reversely translocates along the DNA, threading its 3’ transcribed RNA through the secondary channel and rendering Pol II inactive^50-55^. This observation in Fig. 1d shows that backtracking is the primary cause for stalling under torsion, indicating that anti-backtracking transcription elongation factors, such as TFIIS^52^, may facilitate Pol II transcription under torsion by limiting backtracking or rescuing the backtracked complexes in a way similar to the effect of GreB on *E. coli* RNAP^11^.

Investigating whether TFIIS can facilitate Pol II transcription under torsion and, more broadly, how Pol II works against obstacles requires extensive exploration of broad experimental conditions, which is challenging when measurements must be conducted sequentially on the AOT. The scope of this investigation motivated us to develop an alternative method using magnetic tweezers (MT), which enable parallel measurements of multiple molecules while resolving the dynamics of each molecule^56-59^. Although the MT instrument cannot directly measure torque, the torque in the DNA substrate may be obtained through the measured DNA extension and force using the DNA torsional properties established from the AOT (Supplementary Fig. 4).

We first used the MT assay to determine the stall torque of Pol II in the presence of TFIIS. In this experiment, we anchored Pol II to a magnetic bead and torsionally constrained the downstream DNA to the coverslip surface of a sample chamber (Fig. 2a). Previously, we showed that *E. coli* RNAP stall torque may be obtained via a torque jump method using the AOT^10^. We adapted this method for Pol II stall torque measurements on the MT. We first verified that Pol II was active under an assisting torque by allowing it to transcribe under (-) supercoiling (Methods). Continued transcription converts the (-) supercoiling to (+) supercoiling and buckles the DNA to form a (+) buckled plectoneme at +6 pN·nm torque. We then jumped the torque to +10 pN·nm and determined whether Pol II could continue moving forward (Methods). In the example trace without TFIIS (Fig. 2b), Pol II stalls and backtracks, indicating +10 pN·nm is significant resistance to forward movement. In contrast, in the presence of TFIIS (Fig. 2b), Pol II continues to move forward despite the +10 pN·nm resistant torque. During the forward movement, Pol II pauses between steady movements, but the pauses show minimal backtracking, consistent with the anti-backtracking action of TFIIS.

*In vivo*, the torsional stress in DNA is regulated by topoisomerase activity: high torsional stress when topoisomerase activity is low and low torsional stress when topoisomerase activity is high. Our torque jump experiment mimics this process by examining Pol II activity at different torques. Fig. 2c shows the mean trajectories of Pol II. Under low torque values, Pol II moves forward steadily, showing minimal differences with and without TFIIS. However, when the torque increases to +10 pN·nm, Pol II without TFIIS cannot sustain steady forward movement and backtracks extensively; in contrast, Pol II with TFIIS can still move forward. The torque-velocity relation, characteristic of the chemo-mechanical property of a rotary motor protein^10^, shows that Pol II pause-free velocity decreases with an increase in torque (Fig. 2d). Although TFIIS dramatically increases the overall ability of Pol II to move forward against torsional stress (Fig. 2c), TFIIS does not increase the pause-free velocity, consistent with TFIIS acting only on a paused elongation complex (Fig. 2d).

To determine the stall torque of Pol II, we measured the fraction of Pol II molecules remaining active at different torques (Fig. 2e). We define stall torque as the torque when 50% of the molecule remains active^10^. This analysis shows that Pol II alone can generate +8.6 ± 0.3 pN·nm torque, consistent with values measured using the AOT within the measurement uncertainty. The presence of TFIIS increases Pol II stall torque significantly to +12.8 ± 0.4 pN·nm, representing a 49% enhancement. Thus, by limiting backtracking, TFIIS enables Pol II to work more effectively against torsional stress.

As a comparison, *E. coli* RNA polymerase alone generates ∼ +11 pN·nm torque^10^. So, Pol II generates a torque comparable to *E. coli* RNA polymerase, though slightly smaller. In addition, *E. coli* RNA polymerase’s torsional generation capacity is also enhanced by the anti-backtracking factor GreB to +18.5 pN·nm torque^11^, indicating that the torsional capacity enhancement by anti-backtracking transcription elongation factors may be general to all transcription machineries. Overall, the basal machinery of eukaryotic transcription is a powerful rotary motor whose torsional capacity can be regulated by transcription factors. *In vivo*, many other factors may potentially modulate the torsional capacity of Pol II to facilitate its passage through obstacles such as nucleosomes^60^.

### Transcription through nucleosomes under torsion

In recent decades, through extensive *in vitro* and structural studies, we have gained significant mechanistic insights into the nature of the nucleosome barrier to transcription^52, 61-69^. These studies have focused primarily on transcription through a DNA substrate containing a single nucleosome under no torsional stress. In the cell, Pol II must transcribe through nucleosomes in chromatin while supercoiling DNA. Thus, Pol II simultaneously experiences a physical blockage from the encountered nucleosome and resistance from DNA supercoiling buildup in the chromatin. The mechanistic understanding of this process has remained obscure due to the complexity of this problem and a lack of experimental approaches for investigation.

To tackle this problem, we have developed a real-time, magnetic tweezers-based assay to track the progression of Pol II as it transcribes through nucleosomes in a chromatin fiber (Fig. 3a). The experimental configuration is similar to that of Fig. 2a, except that the downstream template contains either a single nucleosome or a chromatin fiber containing about 50 nucleosomes. To facilitate comparison, the chromatin fiber template is identical to the single-nucleosome template in the number of base pairs, the transcription start-site location, and the sequence leading to and through the first nucleosome (Supplementary Fig. 3b). Control experiments show that the nucleosomes are stable under the experimental conditions used here (Supplementary Fig. 4).

**Figure 3.**
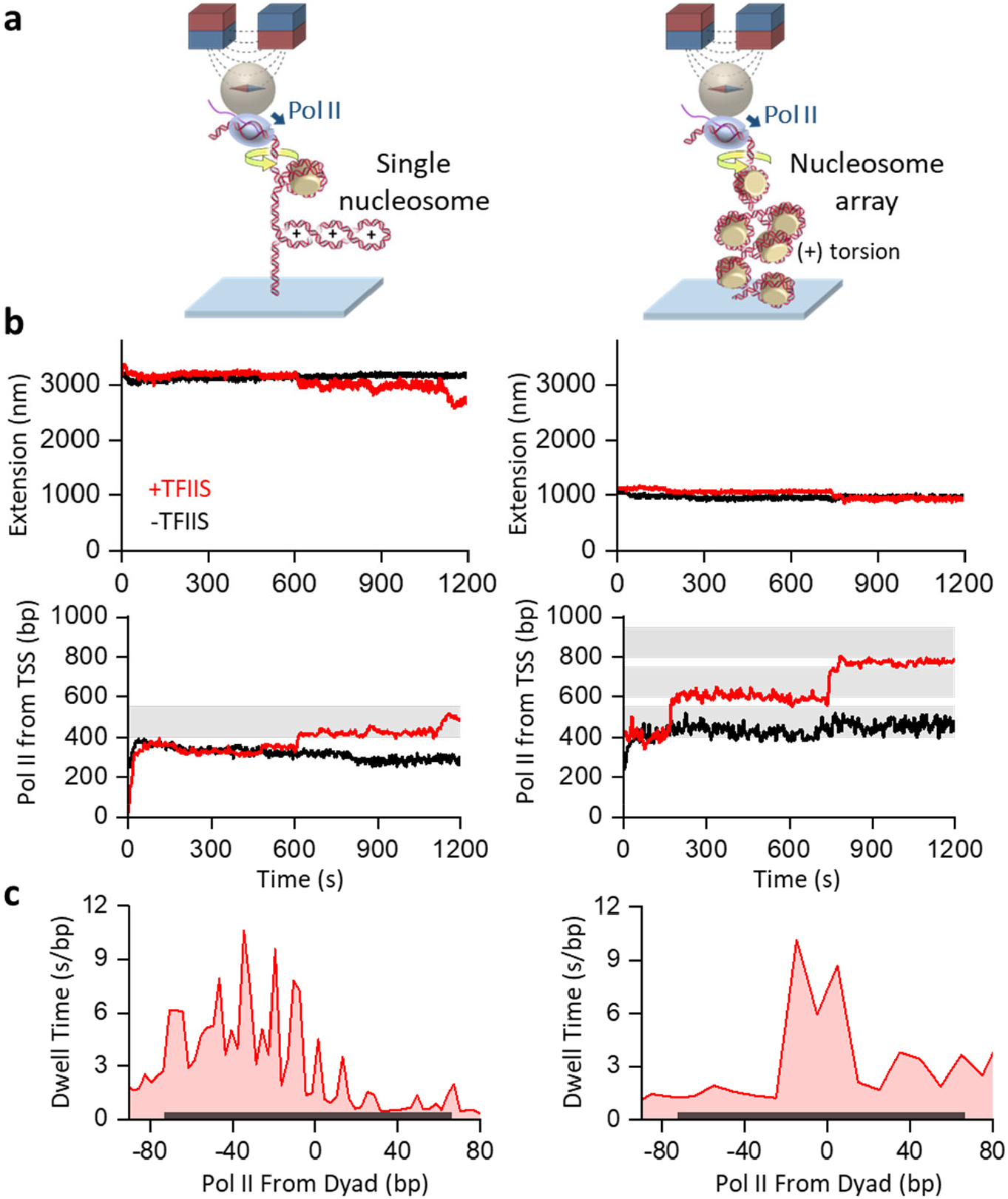
A real-time assay of tracking Pol II transcribing through nucleosomes under torsion via magnetic tweezers. **a**. Experimental configurations. Pol II is torsionally anchored to the surface of a magnetic bead. The DNA downstream of Pol II could contain a single nucleosome (left) or a nucleosome array (right) (Supplementary Fig. 3b) and was torsionally anchored to the surface of a coverslip. Despite having different nucleosomes, the two templates are identical in their overall length and in the sequence up to the first nucleosome encountered. **b**. Representative trajectories of Pol II transcribing through nucleosomes under torsion. The left panels show traces from the template containing a single nucleosome. The right panels show traces from the template containing a nucleosome array. The measured extension is converted to the Pol II position from the TSS for each trace (Methods). Each shaded region represents an expected nucleosome position. **c**. Pol II dwell time distribution at a nucleosome encounter. The dwell time at each position after Pol II encountering a nucleosome is calculated from the Pol II trajectories using a “first-passage” method (Methods). The dark grey shaded region indicates the location of the nucleosome positioning element for the first nucleosome encountered.

Prior to transcription restart, as expected, the measured DNA extension is substantially smaller for the chromatin-fiber template due to the nucleosome compaction of DNA (Fig. 3b). Subsequent transcription leads to a decrease in DNA extension as Pol II introduces (+) supercoils to DNA with its progression. We use the real-time DNA extension to monitor Pol II position along the DNA as Pol II moves forward, (+) supercoils the DNA, and encounters the nucleosome obstacle (Supplementary Fig. 6; Methods).

In the representative traces (Fig. 3b), Pol II pauses when encountering a nucleosome under torsion, and the presence of TFIIS significantly increases the ability of Pol II to move through a nucleosome on both the single-nucleosome and chromatin templates. We examined the dwell time of Pol II when encountering a nucleosome in the presence of TFIIS (Fig. 3b). On the single-nucleosome template, Pol II pauses strongly before reaching the dyad of the nucleosome, with a pausing pattern indicative of periodicity (Fig. 3c left panel). Fourier transformation of this signal reveals a 11 bp periodicity (Supplementary Fig. 7), consistent with Pol II pausing at the strong histone-DNA interactions in a nucleosome, which occur whenever the DNA minor groove faces the histone core octamer surface^70, 71^. This exceptional resolution results from the detection resolution enhancement of buckled DNA: each 10.5 bp of Pol II movement leads to a ∼ 50 nm reduction to the DNA extension. The chromatin-fiber template does not allow this resolution enhancement, but Pol II dwelling around the dyad is still detected (Fig. 3c right panel). For both templates, the pausing patterns bear some resemblance to those previously observed *in vitro*^61, 72, 73^ or *in vivo*^74^, with pausing occurring primarily before and around the dyad. These observations demonstrate that we have established a method to investigate how Pol II transcribes through nucleosomes under torsion.

### Chromatin buffers torsion to facilitate transcription

Intriguingly, the representative traces in Fig. 3b are suggestive that Pol II transcribes through a nucleosome more efficiently on the chromatin template than on the single-nucleosome template. To rigorously evaluate this observation, we pooled many traces and plotted the mean Pol II trajectory at their nucleosome encounter (Fig. 4a, Methods). Indeed, Pol II transcription is more efficient on the chromatin template. In the absence of TFIIS, Pol II stalls extensively when it first encounters the nucleosome on the single-nucleosome template. In contrast, Pol II can significantly invade the nucleosome on the chromatin template. The presence of TFIIS greatly facilitates Pol II passage through a nucleosome on both templates, but transcription remains more efficient on the chromatin template.

It is important to note that for both templates, which are *identical* in length, Pol II transcribes the *same* DNA sequence and generates the *same* number of supercoils when encountering the nucleosome (Supplementary Fig. 3b). Yet, the nucleosome-passage rates significantly differ (Fig. 4b). The only difference between the two configurations is the substrate downstream of the nucleosome that Pol II first encounters: naked DNA for the single-nucleosome template and chromatin fiber for the chromatin template. Our previous studies show that the torsional stiffness of chromatin fiber is about 8 times softer than that of naked DNA^57^. Thus, the torsional mechanical properties of chromatin allow effective buffering of torsion generated from transcription on the chromatin template^57, 75^, facilitating transcription through the nucleosome, while naked DNA of the single-nucleosome template cannot effectively buffer torsion generated from transcription, making torsion a significant obstacle to transcription.

**Figure 4.**
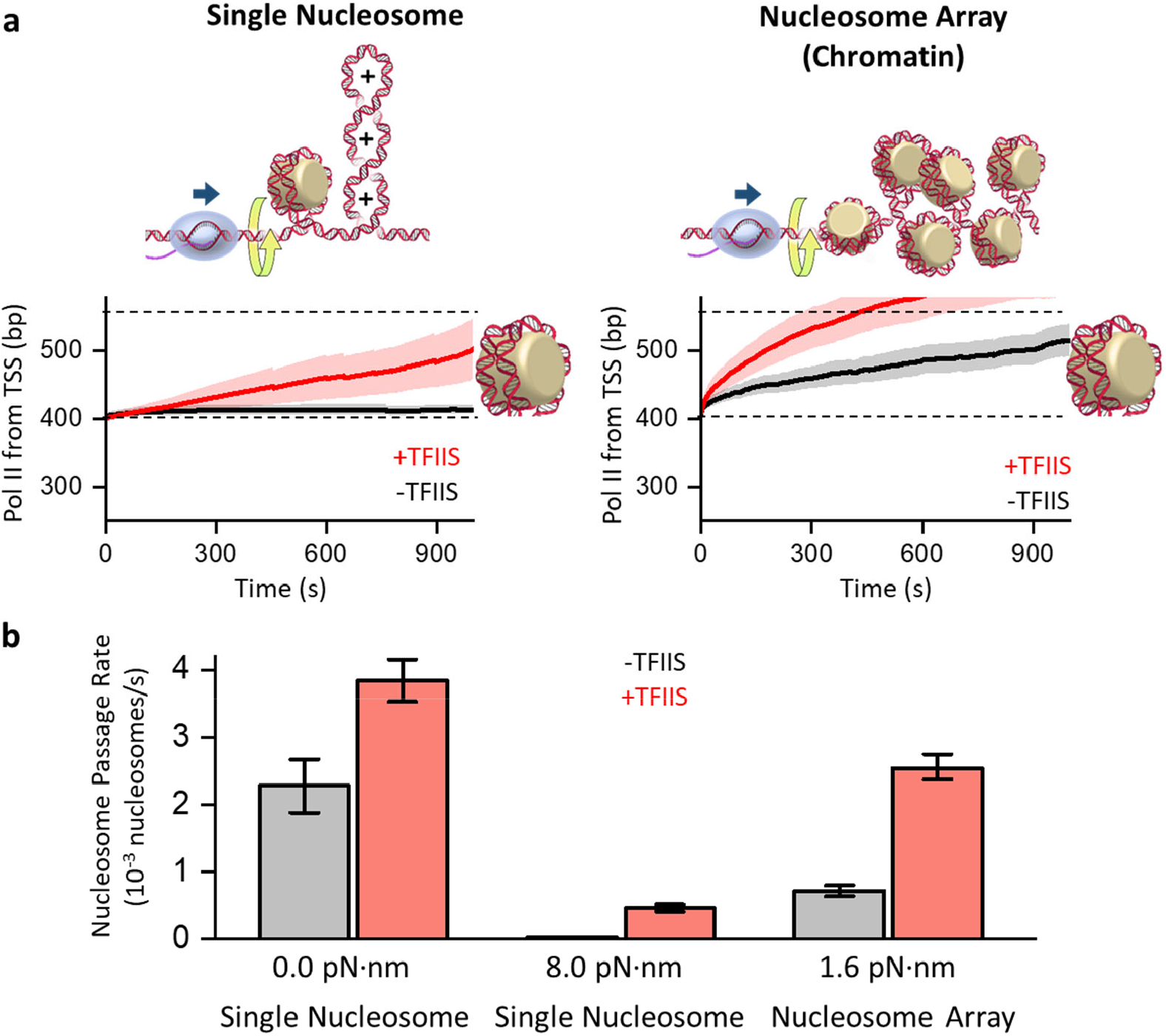
Chromatin buffers torsional stress to facilitate transcription. **a**. Mean trajectories of Pol II transcription through a nucleosome. All traces were aligned when Pol II reached the entry of the 1^st^ nucleosome encountered (*t* = 0), with the shaded regions representing 30% of the standard deviation. The left panel shows data from the single-nucleosome template (left; *N* = 86 for -TFIIS, *N* = 215 for +TFIIS). The right panel shows data from the first nucleosome on the chromatin template (right; *N* = 99 for -TFIIS; *N* = 302 for +TFIIS). **b**. Pol II nucleosome-passage rate. The mean passage rates under torsion were obtained from data shown in **a**, with the torque value being the torque experienced by Pol II when encountering the nucleosome dyad (Methods). The mean passage rates under zero-torsion of the single-nucleosome template were obtained from data shown in Supplementary Fig. 8. The error bars represent the SEMs.

Consistent with this chromatin torsional mechanical model, our measured nucleosome passage rate is much greater on the chromatin template than on the single-nucleosome template. It is 6-fold greater in the presence of TFIIS. Using the torsional mechanical properties of chromatin^57^, we estimate that Pol II experiences only 1.6 pN·nm resisting torque when encountering a nucleosome on the chromatin template but 8.0 pN·nm resisting torque when encountering a nucleosome on the single-nucleosome template (Methods). To further evaluate this possibility, we also examined the nucleosome-passage rate on a single-nucleosome template that is not torsionally constrained so that torsion cannot accumulate on this template (0 pN·nm) (Supplementary Fig. 8). Indeed, the nucleosome-passage rate on this template is significantly increased and even greater than that on the chromatin-fiber template under torsional stress, further demonstrating how torsion can impact the nucleosome-passage rate.

Collectively, our findings demonstrate the effectiveness of chromatin in buffering torsional stress during transcription. Although chromatin has been traditionally viewed as a roadblock to transcription^65, 66^, our results show that by buffering torsional stress, chromatin actually facilitates transcription, revealing a crucial role of chromatin in regulating transcription. Importantly, although DNA supercoiling and torsion are distinct physical quantities, they are often conflated in the literature. Our work shows how torsion can significantly impact transcription even under the *same* degree of supercoiling. Torsion, not DNA supercoiling, dictates the difficulty of Pol II in twisting DNA. The distinction between DNA supercoiling and torsion is critical but historically overlooked. Our data help highlight this distinction.

### Torsion modulates nucleosome passage

Even though chromatin is an effective torsional buffer, continuous transcription eventually generates enough torsion to exceed the buffering capacity of chromatin, limiting Pol II’s ability to transcribe through multiple nucleosomes. Thus, timely torsional relaxation by topoisomerases is crucial to ensure Pol II progression^14, 19-24^.

To investigate how topoisomerase relaxation facilitates transcription through chromatin, we conducted transcription experiments in the presence of yeast topoisomerase II (topo II) (Fig. 5a) using a configuration similar to that shown in Fig. 3a. We used 100 pM of yeast topo II, which can effectively relax torsional stress in a chromatin substrate^57, 58^. However, the Pol II location on DNA can no longer be tracked in real-time by the DNA extension, since the DNA extension can be altered by topo II relaxation or Pol II translocation and thus does not uniquely inform the Pol II location on the template. Instead, we assess the Pol II location using the extension-turns relation of the chromatin substrate downstream of Pol II. Previously, we have characterized how the width and height of the extension-turns relation of chromatin depend on the number of nucleosomes^57, 58^. We now use this characterization to estimate the number of nucleosomes on the remaining chromatin between Pol II and the distal DNA. As Pol II progresses through nucleosomes, the chromatin array ahead of Pol II shortens, leading to a reduced width and height of the extension-turns relation (Fig. 5b). These features provide an estimation for the number of nucleosomes transcribed.

**Figure 5.**
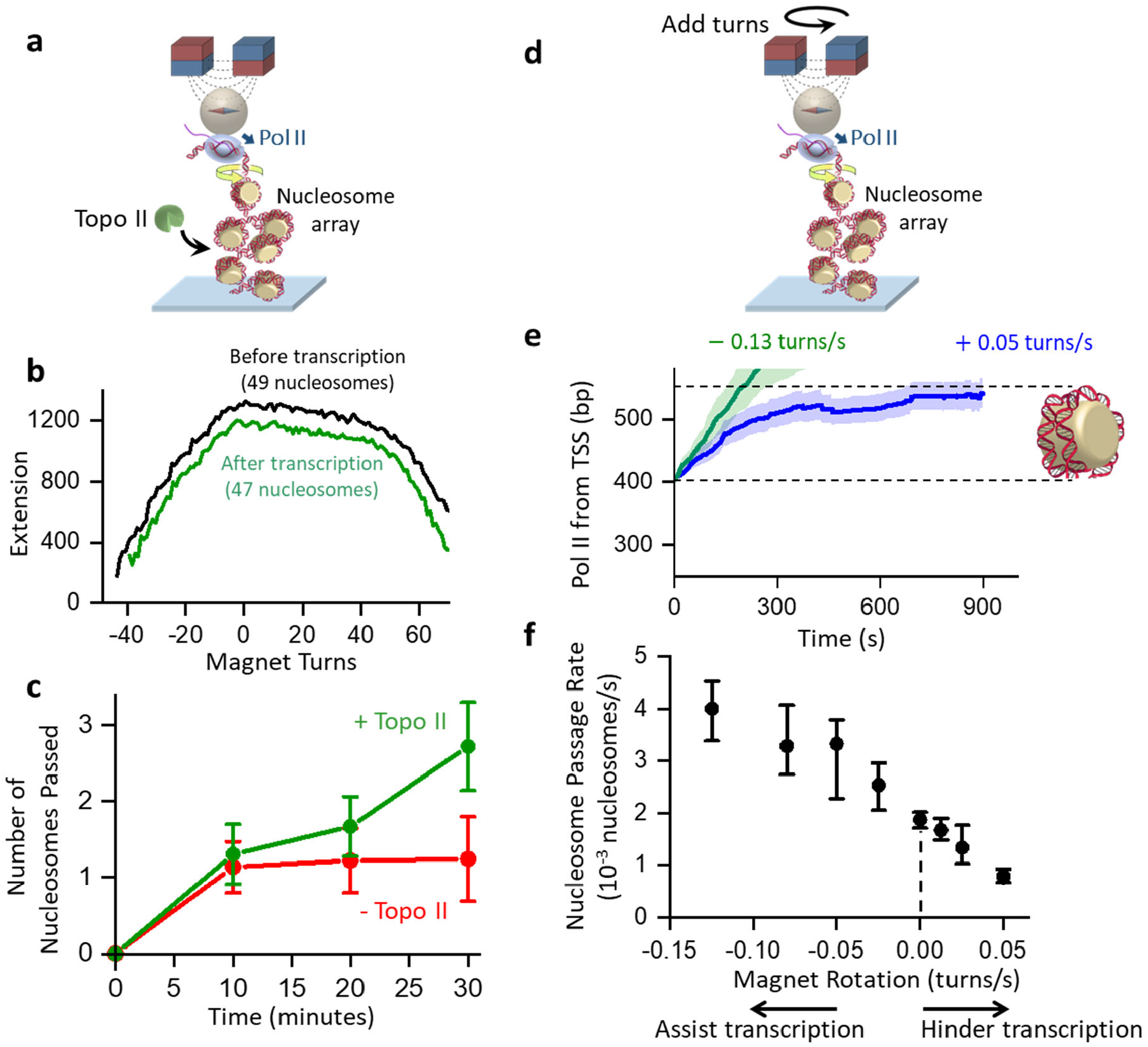
Torsion modulates nucleosome passage. **a**. Experimental configuration of transcription through chromatin in the presence of topo II. This configuration is identical to that used in Fig. 3a (right), except for having topo II present in the assay. All experiments were carried out in the presence of TFIIS. **b**. Explanation of the method used to determine the number of nucleosomes passed. The extension-turns relation is determined by the number of nucleosomes and the DNA length between Pol II and the bead. Both the width and height of the extension-turns curve decrease after transcription. Shown is an example trace where we measured 49 nucleosomes before transcription and 47 nucleosomes after transcription, indicating Pol II having passed 2 nucleosomes during transcription. **c**. The number of nucleosomes Pol II passes through during transcription versus transcription time. The error bars represent the SEM, with each data point collected from *N* = 14-33 traces. **d**. Experimental configuration to modulate torsion using the magnetic bead during transcription through chromatin. This configuration is similar to that used in Fig. 3a (right), except that the magnet bead is rotated at a constant rate. All experiments were carried out in the presence of TFIIS. **e**. Mean trajectories of Pol II transcription through a nucleosome. All traces were aligned when Pol II reached the entry of the 1^st^ nucleosome encountered (*t* = 0), with the shaded regions representing 30% of the standard deviation. Shown are examples of magnet rotation rates at +0.05 turns/s (hindering transcription) (*N* = 30) and −0.13 turns/s (assisting transcription) (*N* = 49). **e**. Pol II nucleosome-passage rates at different magnet rotation rates. The mean rates for the passage of the 1^st^ nucleosome encountered are shown, with the error bars representing the ± SEMs, with each data point collected from *N* = 25-65 traces.

Using this method, we monitored how Pol II transcribes through chromatin as a function of time (Fig. 5c). In the absence of topo II, Pol II significantly slows down after transcribing through one nucleosome, consistent with torsional resistance buildup. In contrast, when topo II is present, Pol II can continue to transcribe through multiple nucleosomes with little indication of being slowed down over time. Thus, the presence of topo II ensures steady transcription.

*In vivo*, Pol II may also encounter other motor proteins that share the same DNA substrate, and Pol II progression is modulated by torsion generated by those processes (Supplementary Fig. 9). For example, when one Pol II trails another Pol II moving co-directionally, the leading Pol II generates (-) supercoiling behind that annihilates the (+) supercoiling in front of the trailing Pol II, reducing the torsional stress of the trailing Pol II. Similarly, when a Pol II encounters another Pol II or a replisome moving head-on, (+) supercoiling accumulates between the two motors, increasing the torsional stress.

To investigate how additional torsional stress impacts transcription, we allowed Pol II to transcribe on the chromatin template while either mechanically unwinding the DNA to reduce torsion or overwinding the DNA to increase the torsion, mimicing the torsional action of other motor proteins. In this experiment, we rotated the magnet at a constant rate while monitoring the progression of Pol II through a nucleosome (Fig. 5d). As expected, unwinding the chromatin substrate facilitates Pol II progression through a nucleosome while overwinding the chromatin substrate hinders its progress (Fig. 5d,e), consistent with a decrease in transcription activity with an increase in the torsional stress.

Taken together, these results further highlight the role of torsion during transcription through nucleosomes and provide additional support for the torsional mechanical model of transcription. We found that topoisomerase relaxation of the torsional stress can significantly facilitate transcription through chromatin, allowing Pol II to progress through multiple nucleosomes without significant hindrance from the torsion buildup. We also show that Pol II passage through a nucleosome can be modulated by additional torsional stress, which could be imposed by other motors on the same substrate. These results demonstrate how torsion can modulate transcription through nucleosomes.

## Discussion

Our findings show that torsion is a strong regulator of transcription through chromatin. We have provided unprecedented information on Pol II transcription dynamics through chromatin under torsion. This work shows a complex interplay of DNA supercoiling generated by Pol II and chromatin’s ability to buffer the resulting torsion. The direct comparison of transcription through a nucleosome on a single-nucleosome template and a chromatin template also calls attention to the crucial distinction between DNA supercoiling and torsion, which have often been conflated in the literature.

We visualized Pol II rotation of DNA, demonstrating the inevitable consequence of torsion associated with transcription (Fig. 1a,b). Although RNAP has primarily been examined as a linear motor that generates force to translocate along the DNA^40, 41, 45^, this work on Pol II, together with previous studies of *E. coli* RNAP^10, 11^, brings to the fore that RNAP is also a rotary motor that generates torques to rotate the DNA. We found that Pol II can generate a torque of about 9 pN·nm that nearly melts the DNA and becomes significantly more powerful in the presence of TFIIS to about 13 pN·nm (Fig. 1; Fig. 2). Thus, Pol II is a potent rotary motor with its torsional capacity modulated by transcription factors. Although this work focuses on the basal transcription machinery with TFIIS, other factors^76-79^ will likely further modulate the torsional capacity of Pol II *in vivo*.

Pol II generated (+) torsion could destabilize nucleosomes ahead of Pol II, since each nucleosome traps about one (-) turn by the left-handed wrapping of DNA around a nucleosome^80, 81^. We previously found that +19 pN·nm can facilitate the dissociation of the H2A/H2B dimers in a nucleosome^82^, suggesting that the torque generated by Pol II may not be sufficient to destabilize nucleosomes significantly. Consistent with this possibility, our control experiment shows that nucleosomes remain stable during the measurement time under a torque similar to that generated by Pol II (Supplementary Fig. 5). Although the Pol II-generated (+) torque is insufficient to destabilize nucleosomes in our experiments, nucleosome remodelers and chaperons could destabilize the nucleosomes *in vivo*, along with the assistance of torsion, to allow Pol II passage^83-89^.

Our data demonstrate that chromatin can effectively buffer torsional stress generated by transcription, allowing Pol II to progress more efficiently through nucleosomes, a feat unattainable on naked DNA (Fig. 3; Fig.4). Although chromatin has been previously proposed to serve as a torsion buffer ^57, 90, 91^, our data represent the first experimental evidence to support this hypothesis in the context of transcription and demonstrate the role of chromatin beyond being just an obstacle. Interestingly, genomics studies show that nucleosomes are present within gene bodies at high occupancy, even for highly transcribed genes^92, 93^, and deletion of factors that preserve the nucleosome integrity on the chromatin could result in a down-regulation of genic transcription^85^. Thus, chromatin may be a necessary substrate constituent to limit torsion built up during active transcription.

We also show that yeast topo II can greatly facilitate transcription through nucleosomes (Fig. 5a,b,c). Previously, we found yeast topo II to be highly processive and efficient on a chromatin substrate^58^. This may be explained by topo II’s preference for binding DNA crossings, which are readily available at the juxtaposing entrance and exit points of a nucleosome^70, 94^. *In vivo*, other topoisomerases (e.g., topo I) and nearby motor proteins can further modulate torsional stress^17, 95^. We show that transcription through chromatin can be upregulated by (-) torsion or down-regulated by (+) torsion.

While the torque capacity of Pol II may facilitate the ability of the enzyme to transcribe against torsional resistance, this property may also exacerbate conflicts with other motor proteins. For example, when Pol II encounters a replisome head-on, (+) torsion accumulates between Pol II and the replisome, which could occur well before Pol II physically encounters the replisome since torsion can act over a distance. Previous studies support a model in which (+) torsion accumulation is the culprit for replication stress, leading to replication fork stalling and disassembly^27-29^. Although the replisome often ultimately wins the conflict^95-97^, Pol II’s ability to move forward against strong torsional stress will allow a greater (+) torsional buildup between the two motors that leads to greater fork instability, which is detrimental to genome integrity.

Despite chromatin being traditionally viewed as a hindrance to fundamental DNA processes, our findings challenge this notion by demonstrating the role of chromatin in promoting transcription under torsion. Previously, we provided evidence that chromatin facilitates replication by partitioning supercoiling behind the replication fork, simplifying the topology for subsequent chromosome segregation^57^. This work extends this idea in the context of transcription. These studies reveal the crucial role of chromatin’s torsional mechanical properties in fundamental DNA processes.

## ACKNOWLEDGEMENTS

We thank members of the Wang Laboratory for helpful discussion and comments. This work is supported by the National Institutes of Health grants R01GM136894 (to M.D.W.) and T32GM008267 (to M.D.W.). M.D.W. is a Howard Hughes Medical Institute investigator. This work has been performed in part at the Cornell NanoScale Facility, a member of the National Nanotechnology Coordinated Infrastructure (NNCI), which is supported by the National Science Foundation (Grant NNCI-2025233).

## AUTHOR CONTRIBUTIONS

J.Q., J.T.I, and M.D.W. designed single-molecule assays. J.Q. and T.M.K. prepared DNA substrates. L.L. and M.K. purified and characterized Pol II and TFIIS. R.M.F. purified and characterized histones. J.J. and J.M.B. purified and characterized yeast topo II. J.Q. and S.Z. performed single-molecule experiments. Y.H. fabricated and calibrated quartz cylinders. J.Q. and J.T.I. analyzed data. C.T. performed preliminary experiments during the early stage of the project. D.G. created the yeast strain for Pol II purification. M.D.W. wrote the initial draft. All authors contributed to the manuscript revision. M.D.W. supervised the project.

## Code availability

Data analysis routines used to process and generate plots are available in Github: https://github.com/WangLabCornell/Qian_et_al.

## DATA AND MATERIAL AVAILABILITY

All data are available in the main text or the supplementary materials.

