## Supplementary Materials for "Chromatin Buffers Torsional Stress During Transcription"

**Protein expression and purification**

Yeast Pol II containing a 6×His-tag and a biotin tag on both Rpb1 and Rpb3 subunits was expressed and purified from *S. cerevisiae*[1]. In brief, cell pellets were lysed and went through a HisTrap HP column (Cytiva 17524801), a heparin column (Cytiva 17040703) and a Mono Q column 10/100 (Cytiva 17516701). The resulting product was then biotinylated by GST-BirA and the reaction mixture was slowly loaded to a Mono Q column 5/50 (Cytiva 17516601). The eluted biotinylated Pol II was then checked by SDS-PAGE 4-12% Bis-Tris SDS gels in 1X MES SDS buffer, concentrated, and flash-frozen for storage.

*S. cerevisiae* recombinant TFIIS was expressed in *BL21 (DE3) pET15b PPR1* (69450-M; Novagen-sigma)[2]. Cell pellets harvested by centrifugation were sonicated using a microprobe tip and Sonicator-Ultrasonic Processor VHX 750 watt (model GEX 750; PG Scientific). Each chromatography step was checked by SDS-PAGE with 12% Bis-Tris SDS gels in 1X MES SDS buffer. The centrifugated lysate was loaded/passed through a HisTrap FF Crude (Cytiva 17528601) and later was loaded to a Mono S column 10/100 GL (Cytiva 17516901). The protein eluted from the column was diluted and reloaded to a HisTrap HP (Cytiva 29051021) column and a Mono S column 5/50 GL (Cytiva 17516801) to produce 6×His-tagged protein. The protein was then checked by SDS-PAGE, concentrated, and flash-frozen for storage.

Human Hela histones were purified from nuclear pellets of HeLa-S3 (National Cell Culture Center HA.48) cells, followed by a hydroxyapatite Bio-gel HTP gel slurry (Bio-Rad Laboratories) purification[3-5]. The protein was then concentrated and flash-frozen for storage.

Yeast topoisomerase II (topo II) was expressed in *S. cerevisiae*[6-8]. In brief, Cell pellets harvested by centrifugation were lysed, and went through a HisTrap HP column (Cytiva 17524802) and a HiTrap SP HP column (GE 17115201). The protein eluted from the column was concentrated and incubated with TEV protease (QB3 Macro Lab) overnight, removing the 6×His-tag to produce tag-free proteins. The protein was further purified using a HisTrap HP column (Cytiva 17524802) and a Sephacryl column (GE Healthcare). The protein was then concentrated and flash-frozen for storage.

**DNA template construction**

The 'gapped' DNA construct (Supplementary Fig. 2) was formed on a 0.6 kb DNA PCR amplified from pMDW139 (see Supplementary Table 1 for primer sequences). Two nicks on the non-template strand (NTS) were introduced by Nt.BbvCI (NEB R0632). The sample was then melted and re-annealed with the addition of 40X amount of a DNA oligo with an identical sequence to the template strand (TS) (ARP78 TS, Supplementary Table 1) and went through spin column purification (Select-a-Size DNA Clean & Concentrator, Zymo Research D4080), resulting in a 76 nt gap on the non-template strand. The sample was further digested with BstXI (NEB R0113) to create a unique overhang to ligate to the downstream template.

A second 'gapped' DNA construct of 133 bp used for the Pol II upstream torque was formed by annealing three DNA oligos: Upstream NTS, Downstream NTS and Short TS (see Supplementary Table 1 for sequences). The resulting DNA segment contains two unique overhangs at the template ends.

The 5.7 kb DNA construct (Supplementary Fig. 3a) for Pol II rotation tracking and downstream stall torque measurement is composed of a 4.6 kb center segment (~50% GC content) flanked by a 0.6 kb transcription elongation complex (TEC) segment and a 0.5 kb multi-digoxigenin-labeled tethering adapter. The 4.6 kb center segment was PCR amplified from pRL54 (see Supplementary Table 1 for primer sequences) with LongAmp Taq (NEB M0534) and digested with BstXI (NEB R0113) and BsaI-HFv2 (NEB R3733) to produce unique overhangs.

The 11.6 kb DNA construct (Supplementary Fig. 3a) for Pol II upstream stall torque measurement is composed of a 0.5 kb multi-digoxigenin-labeled tethering adapter, a 6.5 kb upstream segment (~50% GC content), a 133 bp TEC segment and a 4.6 kb center segment. The 6.5 kb upstream segment was PCR amplified from pMDW133 (see Supplementary Table 1 for primer sequences) and digested with BssSI-v2 (NEB R0680) and PpuMI (NEB R0506) to produce unique overhangs.

The 601 sequence[9] was used as the nucleosome positioning element (NPE) for both single nucleosome and nucleosome array templates. The 13.8 kb single-nucleosome template (Supplementary Fig. 3b) is composed of a 0.6 kb TEC segment, a 0.3 kb nucleosome segment containing a single NPE, a 12.3 kb center segment, and a 0.5 kb multi-digoxigenin-labeled tethering adaptor[6, 10]. The reverse PCR primer of the 0.6 kb TEC segment was modified (0.6 kb gapped PCR R Nuc, Supplementary Table 1) to produce the correct overhang for downstream ligation. The 0.3 kb NPE segment was PCR amplified from pMDW2 (see Supplementary Table 1 for primer sequences). The product was then digested with AlwNI (NEB R0514) and BbsI-HF (NEB R3539) to generate overhangs for further ligation. The 12.3 kb center segment was PCR amplified with Phusion (NEB M0530) from λ-DNA (NEB N3011, see Supplementary Table 1 for primer sequences) and digested with BstXI (NEB R0113) and BsaI-HFv2 (NEB R3733) to generate unique overhangs for further ligation.

The 13.8 kb nucleosome array template (Supplementary Fig. 3b) is composed of a 0.6 kb TEC segment, a 12.7 kb 64-mer segment containing 64 tandem repeats of 197 bp (147 bp NPE +

50 bp linker DNA), and a 0.5 kb multi-digoxigenin-labeled tethering adaptor. The 64-mer segment was digested with FastDigest BglI (Thermo Scientific FD0074) and FastDigest BstXI (Thermo Scientific FD1024) from pMDW72 and went through gel extraction[6, 11].

**Pol II EC formation**

Pol II was directly assembled into a transcription elongation complex (TEC)[11-13]. A 'gapped' DNA construct was formed with a 76 nt gap on the non-template strand (NTS) (Supplementary Fig. 2). A 14 nt RNA (RNA 14, Supplementary Table 1) was annealed to the gapped region, forming an RNA-DNA hybrid. Pol II was added to the hybrid, followed by the addition of a 76 nt non-template strand DNA oligo (ARP77 NTS, Supplementary Table 1) complementary to the gapped sequence. The nicks at the two ends of the gap were ligated by T4 DNA Ligase (NEB M0202).

**Nucleosome assembly**

Nucleosomes were assembled using the purified HeLa histones onto DNA constructs via salt dialysis[3-5]. The NaCl concentration during the dialysis was decreased from 2.0 M to 0. The quality of the assembly was initially assessed by gel electrophoresis. For single nucleosome constructs, the quality of the assembled nucleosome was later rigorously assessed with an unzipping assay using optical tweezers[14]. For nucleosome array constructs, the quality and occupancy of the assembled nucleosomes were later rigorously assessed with a stretching assay using optical tweezers[3-6].

**Experimental conditions**

The assembled Pol II transcription elongation complex was stored in TEC storage buffer (25 mM Tris-HCl pH 8, 150 mM KCl, 1 mM $MgCl_2$, 10 µM $ZnSO_4$, 1.5 mg/mL β-casein, 1 mM DTT, 2 mM TCEP, 3% glycerol). Transcription was resumed by the addition of 1 mM of each ribonucleoside triphosphate (NTPs, Roche 11277057001) in the transcription buffer (25 mM Tris-HCl pH 8, 150 mM KCl, 5 mM $MgCl_2$, 1 mM NTPs, 10 µM $ZnSO_4$, 1.5 mg/mL β-casein, 1 mM DTT, 2 mM TCEP, 3% glycerol). For experiments requiring TFIIS, 500 nM TFIIS was added to the transcription buffer. For experiments requiring yeast topo II, 100 pM topo II was added to the transcription buffer. All experiments were performed at room temperature.

**Single-molecule assay on angular optical trap (AOT)**

Experiments described in Fig. 1 using the AOT[15, 16] were performed with the biotinylated Pol II being torsionally anchored on the surface of the chamber via biotin-streptavidin interaction, and the downstream multi-digoxigenin-labeled tethering adaptor being torsionally anchored to the bottom surface of an anti-dig coated cylinders quartz cylinder. To enable this measurement, we modified our previous methods for nanofabrication of the quartz cylinders[6, 10, 17] before subsequently coating the cylinders with anti-digoxigenin (Supplementary Fig. 1).

These experiments were performed in nitrocellulose-coated sample chambers[6-8]. The surfaces were functionalized with biotinylated bovine serum albumin (Thermo Scientific 29130) and streptavidin (Agilent SA10-10), then passivated with β-casein (Sigma C6905). DNA was tethered to the surface via Pol II before the quartz cylinders were introduced to the chamber. The laser power entering the objective during all AOT experiments was kept at 20 mW to minimize the possibility of photo-damage.

During the Pol II rotation experiments (Fig. 1b), transcription was resumed by introducing 1 mM NTPs in the transcription buffer. The DNA construct was mechanically unwound until (−) supercoiled plectonemes were formed under a force clamp of 0.3 pN by modulating the trap height. This step encourages Pol II to resume transcription and identifies active Pol II tethers. Continued Pol II transcription neutralized the downstream (−) supercoiling and then formed (+) plectonemes. After Pol II transcription formed (+) plectonemes, a force clamp of 0.2 pN was applied by mechanical rotation of the cylinder while the extension of the tether was kept constant. Thus, the rotation of the cylinder directly reflects the turns introduced by Pol II transcription. Because this treatment does not take into account the contour length shortening, the measured rotation rate is accurate within ~3%.

During the Pol II stall torque measurement (Fig. 1c), transcription was resumed as above. After Pol II transcription buckled the DNA while working against a resisting torque, the force clamp was turned off, and the trap height was held constant. The cylinder’s angular orientation was also held constant to allow Pol II accumulation of supercoiling in the DNA. As Pol II transcribed, the force and the corresponding buckling torque (Supplementary Fig. 4) increased until Pol II stalled. The stall torque for each trace is defined as the maximum torque measured within 120 s after the start of stalling. Pol II position along the DNA during the measurement was determined based on the torsional mechanics of the remaining DNA[18, 19].

**Single-molecule assay on magnetic tweezers (MT)**

Experiments described in Figs. 2-5 were performed with the biotinylated Pol II being torsionally anchored on the surface of a streptavidin-coated paramagnetic bead, and the downstream multi-digoxigenin-labeled tethering adaptor being torsionally anchored on the surface of the chamber. Sample chambers were assembled as described above for AOT experiments. The surfaces of the assembled sample chambers were functionalized with anti-digoxigenin (Vector Labs MB-7000), and passivated with β-casein (Sigma C6905). About 20 pM DNA constructs were introduced into the chamber, followed by incubation with magnetic beads (Invitrogen 65601).

Magnetic tweezers (MT) experiments were performed on a home-built instrument[6-8], which permits simultaneous measurements of multiple molecules under a constant force. In each sample chamber, the extensions of ~50 actively transcribing Pol II were simultaneously measured to determine the position of the Pol II. Although the MT instrument cannot directly measure the torque, the torque can be inferred from the force after the DNA construct has buckled to form plectonemes using the well-established relation between force and torque in plectonemic DNA[6, 18, 19] (Supplementary Fig. 4).

In the torque jump assay (Fig. 2a), after the introduction of 1 mM NTPs, the DNA tethers were (−) supercoiled by adding −17 turns under 0.3 pN force to encourage Pol II to resume transcription and identify active Pol II tethers. Pol II was allowed to transcribe for 15 – 40 s, neutralizing the downstream (−) supercoiling and creating (+) plectonemes, corresponding to $\tau_{\text{low}}$ = 6 pN·nm. The torque was then rapidly increased from $\tau_{\text{low}}$ to $\tau_{\text{jump}}$, where we observed the behavior of transcription elongation for the next 15 minutes. For $\tau_{\text{jump}}$ = 4 pN·nm, $\tau_{\text{low}}$ = 4 pN·nm was used. To determine the active fraction of Pol II $f_{\text{active}}$, we considered traces where Pol II was actively transcribing under target torque $\tau_{\text{jump}}$, along with traces where Pol II was not transcribing under $\tau_{\text{jump}}$ but was actively transcribing right before the torque increase. The $f_{\text{active}}$ versus $\tau_{\text{jump}}$ data were fit with:

$$f_{\text{active}} = \frac{1}{1 + \exp\left(\frac{\tau_{\text{jump}} - \tau_{\text{c}}}{\tau_0}\right)}$$

which determined the values of $\tau_c$ and $\tau_0$. The critical torque $\tau_c$, at which $f_{\text{active}}$ = 0.5, provides a measure of the stall torque[18].

In assays probing Pol II transcription through nucleosome substrates (Fig. 3 and Fig. 4), the position of the Pol II was monitored under 0.5 pN. Before starting transcription, the initial extension-turns curve of a tether was measured. Immediately after the introduction of 1 mM NTPs and 500 nM TFIIS (if any), the magnet was wound by –18 turns to encourage Pol II to resume transcription and identify active Pol II tethers. The extension signal was then monitored for 20 minutes (Fig. 3b), during which Pol II transcription neutralized the downstream (−) supercoiling, formed (+) plectonemes, and then encountered the nucleosome (or nucleosome array). Subsequently, the final (post-transcription) extension-turns curve was measured at the end of the 20 minutes.

For the topoisomerase assay (Fig. 5a,b,c), topo activity was inhibited by flushing the chamber with TEC storage buffer just before the measurement of the post-transcription extension-turns curve.

For torsional modulation assays (Fig. 5d,e,f), the magnet was mechanically wound at the indicated rate during the 20-minute observation.

**Pol II position during transcription through a nucleosome**

For MT experiments, the position of Pol II on the template is determined by the contour length of DNA between the Pol II and the magnetic bead. The contour length of DNA must be determined from the measured quantities of force, extension, and number of turns mechanically introduced. This conversion can be made using the mechanical properties of the supercoiled DNA and chromatin[6, 8] with the two constraints. The first is that Pol II tightly tracks the DNA helical grove, resulting in Pol II rotating DNA by one turn for each 10.5 bp transcribed[20]. The second one is that each nucleosome constraints −1 turn in the 147 bp DNA[21, 22].

Given these constraints, DNA extension during transcription can be converted to the Pol II position along the template as long as the extension versus magnet turns curve (extension-turns curve) before transcription is measured (Supplementary Fig. 6). This curve was measured for each tether. At a given Pol II position on the template, the expected extension-turns curve is scaled from this initial extension-turns curve based on the DNA and nucleosomes in the remaining template. If a Pol II position is associated with DNA in a nucleosome, the expected extension-turns curve is obtained via a linear interpolation of the curve when Pol II encounters the nucleosome and the curve when Pol II exits the nucleosome. Therefore, as Pol II continues to transcribe, the expected extension-turns curve decreases in the overall height and overall width. Importantly, the curve also shifts to the left due to the accumulation of Pol II generated (+) supercoiling in the DNA.

**Pol II dwell time encountering a nucleosome**

We aim to find Pol II dwell time when Pol II encounters a nucleosome using the measured Pol II position-time trajectories. Pol II backtracks when experiencing high resisting torque (Fig. 2b) and encountering nucleosomes (Fig. 3b). The extent of such reverse motion occurs at a wide range, adding noise to the signal. Thus, we employed a “first passage” method for processing Pol II trajectories prior to generating the average Pol II position and dwell-time histograms (Fig. 3c; Fig. 4; Fig. 5e,f). For every measured Pol II transcribed distance trajectory $x(t)$, we used the maximum distance transcribed $X(t) = \max\{x(t_0)|_{t_0 \leq t}\}$. This method mimics the transcript length observed by gel electrophoresis, especially in the absence of TFIIS.

**Mean trajectory of Pol II**

The mean trajectory of Pol II is determined by pooling many individual traces of Pol II (Fig. 2c, Fig. 4a and Fig. 5e). For an ensemble of traces with individual Pol II trajectories $\{x_i(t)\}_{i=1,2,3\ldots}$, the mean trajectory is given by $\langle x \rangle(t) = \langle x_i(t) \rangle_{i=1,2,3\ldots}$. For nucleosome encounter (Fig. 4a and Fig. 5e), all individual traces $x_i(t)$ were aligned so that Pol II first encounters the nucleosome at $t = 0$.

When no torsion was present on a single nucleosome template (Supplementary Fig. 8), once Pol II passed through the only nucleosome on the template and entered the downstream naked DNA region, it translocated at a significantly greater velocity compared to in the nucleosome region. In this case, the trajectories of individual traces downstream of the nucleosome would greatly impact the averaged trajectory. As such, we applied a "periodic boundary condition" to better mimic a chromatin substrate. For a single Pol II position-time trajectory $x(t)$ (aligned so that Pol II first reaches the nucleosome at $t = 0$) where Pol II exits the first 197 bp (147 bp NPE + 50 bp downstream linker) at $t_\mathrm{e}$, the "periodic boundary condition" trajectory is given by:

$$x_{\mathrm{PBC}}(t) = x(t - \lfloor t/t_\mathrm{e} \rfloor t_\mathrm{e})$$

The mean trajectory is then determined from these "periodic boundary condition" trajectories. This "periodic boundary condition" was only applied to averaged trajectories shown in Supplementary Fig. 8.

**Nucleosome passage rate**

Nucleosome passage rate was introduced to compare Pol II's ability to navigate through the nucleosome roadblock. The nucleosome passage rate is the inverse of the dwell time of Pol II inside the NPE sequence, determined from the mean trajectory. For conditions where Pol II mean trajectory did not completely pass through the NPE sequence during the 20-minutes observation window (e.g., Fig. 4a left panel), we approximated the dwell time by applying a linear fit to the mean trajectory.

**Estimate the torque during Pol II passing through a nucleosome**

When Pol II encounters a nucleosome on a single nucleosome template (Fig. 3a left panel), plectonemes form on the downstream naked DNA template. We used the buckling torque of naked DNA (8.0 pN·nm under 0.5 pN) (Supplementary Fig. 4) as an estimate of the resisting torque Pol II experienced when encountering nucleosome. When Pol II encounters a nucleosome on a chromatin template (Fig. 3a right panel), the resisting torque is determined by the supercoiling state of the downstream nucleosome array template[6]. We use the supercoiling state of the downstream template while Pol II is at the dyad (Pol II elongating +550 bp from the TSS, twisting the downstream DNA for +52 turns in addition to the −18 turns applied by the magnet), estimating the resisting torque on the downstream nucleosome array template to be 1.6 pN·nm.

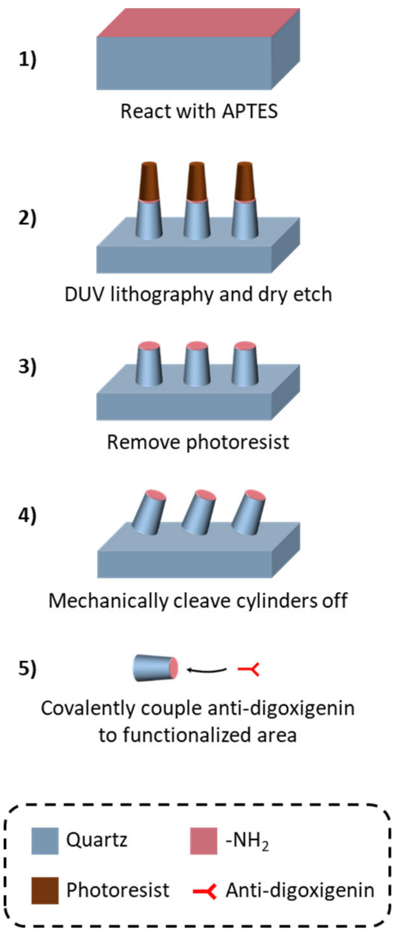


**Supplementary Figure 1**. Fabrication of anti-digoxigenin-coated quartz cylinders.

The nanofabrication process of the quartz cylinders is modified based on a previously established protocol[6, 10, 17]. We first reacted the quartz wafer (Precision Micro Optics, PSQB-131332) with (3-aminopropyl) triethoxysilane (Sigma-Aldrich, 440140) solution. We then dry etched cylinders via a deep ultraviolet (DUV) lithography process. After mechanically cleaving the cylinders off from the wafer, we covalently coupled anti-digoxigenin (Roche, 11333089001) with a linker of glutaraldehyde (Sigma, G5882). These cylinders enable binding of multi-digoxigenin DNA anchors to its functionalized surface in a constrained configuration for DNA torsional studies.

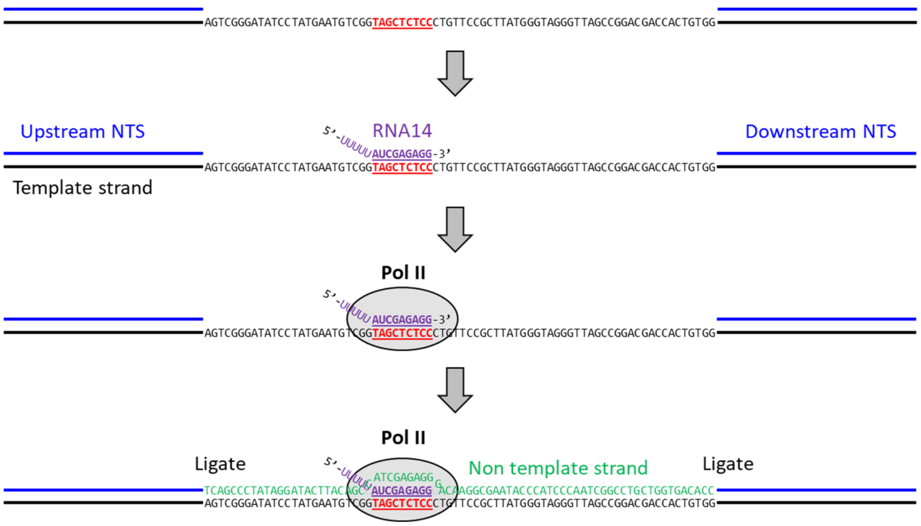


**Supplementary Figure 2:** Pol II elongation complex formation.

A ‘gapped’ template for transcription elongation complex formation contains a 76 nt ‘gapped’ region lacking the non-template strand (the template strand in black; non-template strand in blue; the initial bubble region red and underlined), flanked by dsDNA. A 14 nt RNA oligo (purple) with 9 nt complementary to the template strand is annealed to the ‘gapped’ template. After Pol II binds to the RNA-DNA hybrid, a 76 nt non-template strand DNA oligo (green) with fully complementary sequence to the ‘gapped’ region is added. The nicks at both ends of the ‘gapped’ region are ligated.

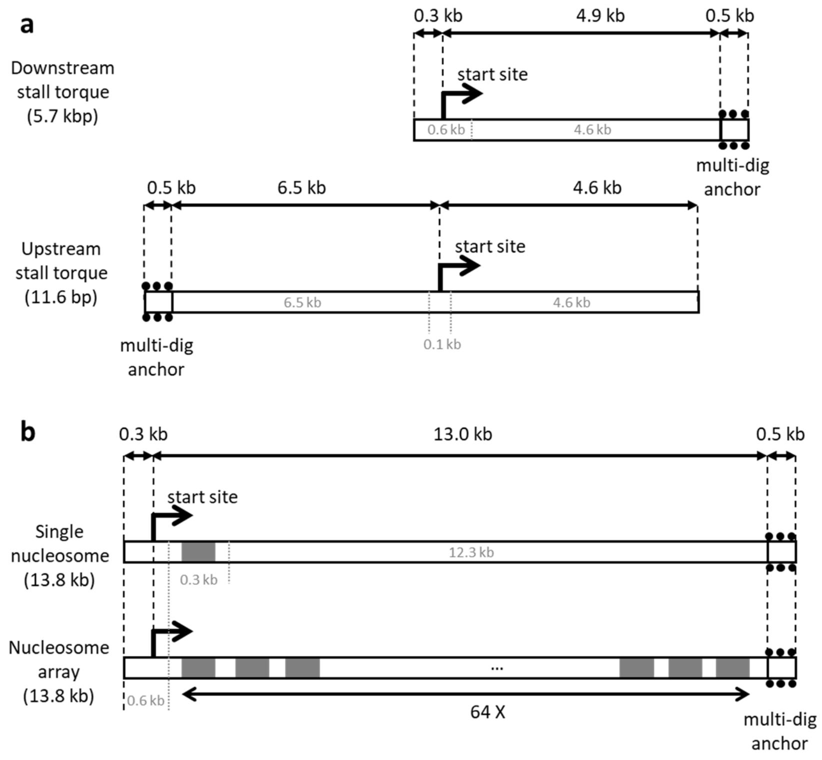


**Supplementary Figure 3:** Single-molecule template design.

**a**. The DNA templates used for the Pol II rotation and stalling experiments.

**b**. The DNA templates used for transcription through nucleosomes experiments. Each 601nucleosome positioning element (NPE) is indicated as a grey box. The first nucleosome encountered is +403 bp from the TSS. The nucleosome array template has a 197 bp (147 bp 601 NPE + 50 bp linker DNA) × 64 tandem repeats. Both the single nucleosome and nucleosome array template are identical in total length and have the same sequence from the upstream end of the template (-313 bp from TSS) to the exit of the first NPE (+550 bp from TSS).

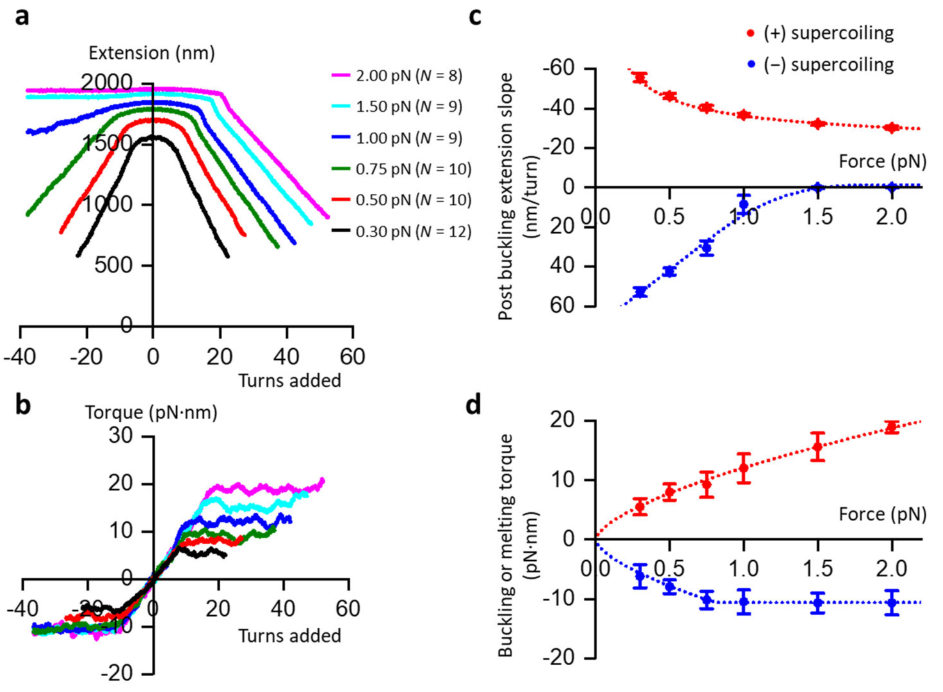


**Supplementary Figure 4.** Torsional properties of naked DNA substrate. A 6.5 kb DNA[8] was used to obtain the torsional properties of naked DNA in the transcription buffer using the AOT.

**a,b**. Extension-turns curves and torque-turns curves under different forces.

**c**. Post-buckling or melting extension slope versus force. The values were obtained by performing a linear fit to the post-buckling or melting region of the extension-turns curve with error bars representing fitting uncertainties.

**d**. Post-buckling or melting torque versus force. The values were obtained by performing a horizontal line fit to the post-buckling or melting region of the torque-turns curve with error bars representing fitting uncertainties.

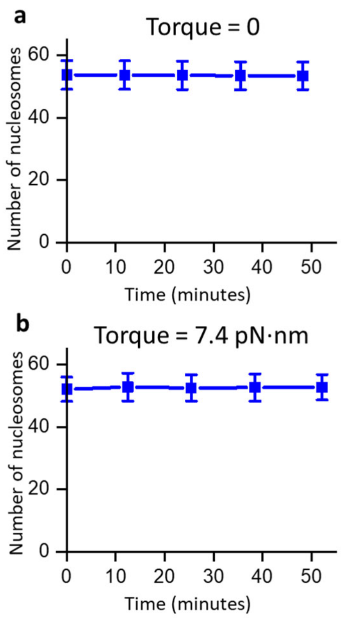


**Supplementary Figure 5.** Nucleosome array stability over time. To assess the stability of the nucleosome arrays, we measured the number of nucleosomes in the array over time in a transcription buffer without transcription.

**a**. Measurements under no torque (*N* = 48). Error bars represent standard deviation of the mean.

**b**. Measurements under 7.4 pN·nm torque (*N* = 26). Error bars represent standard deviation of the mean.

No significant change in the number of nucleosomes over the experimental time window was observed.

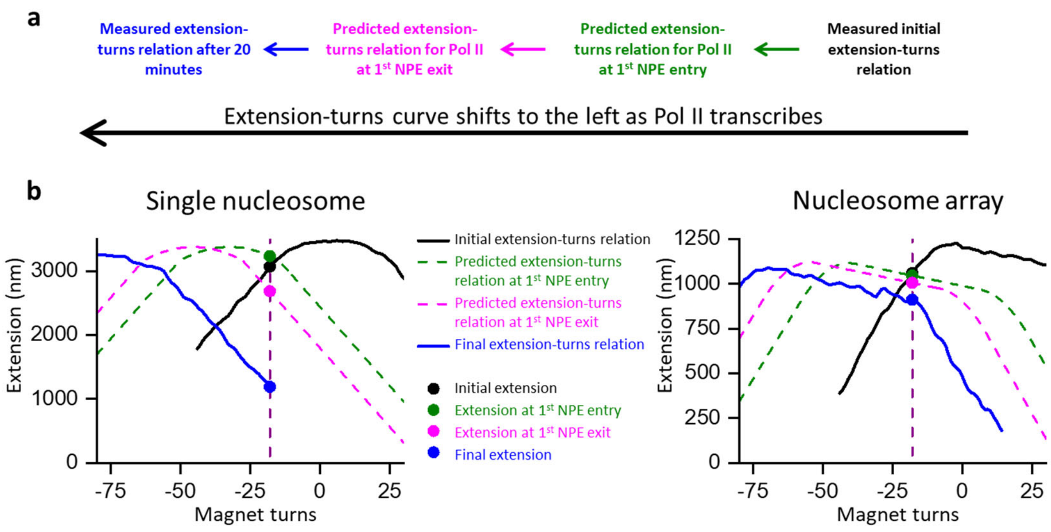


**Supplementary Figure 6.** Pol II position determination.

**a**. The position of Pol II on the DNA template during transcription is determined from the measured extension using the predicted extension-turns curve as a function of Pol II position.

**b**. To demonstrate how this method works, we show examples of how the extension-turns curve changes as Pol II transcribes along the DNA template on a single nucleosome template (left) or on a nucleosome array template (right). In each example, the initial extension-turns curve (black) is measured prior to Pol II transcription. Subsequent Pol II transcription decreases the DNA contour length and increases the DNA linking number density in the downstream DNA segment, resulting in scaling and a shift of the predicted extension-turns curve. The measured extension represents a single point on the predicted extension-turns curve and provides the position of Pol II. The final position of Pol II is further confirmed by the measured extension-turns curve at the end of an experiment.

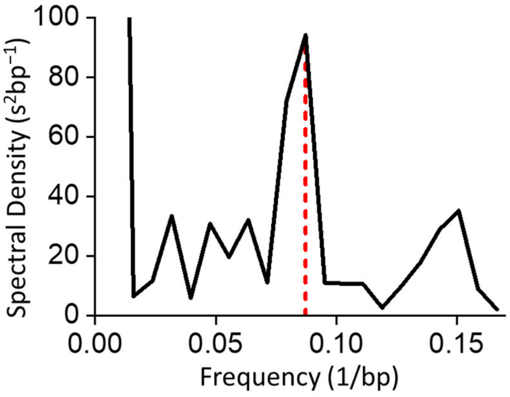


**Supplementary Figure 7.** Power spectral density of Pol II dwell time histogram when encountering the single nucleosome on the single-nucleosome template. The spectra density was obtained from the Fourier transform of data shown in Fig. 3c (left). It reveals a ~ 11 bp periodicity, which agrees well with the helical pitch of dsDNA.

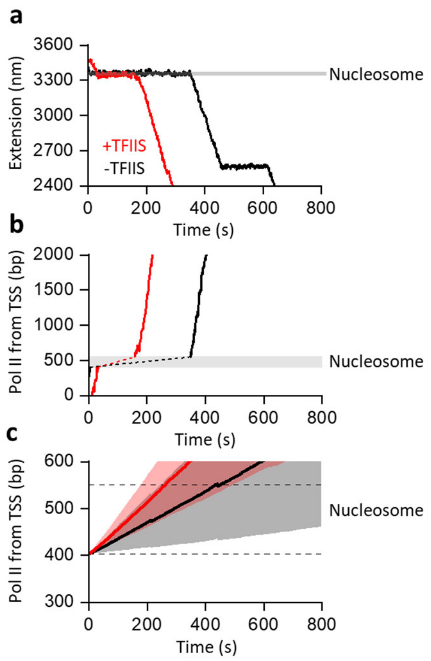


**Supplementary Figure 8.** Pol II transcribing through a single nucleosome under zero torsion.

**a.** Representative trajectories of Pol II transcribing through a single nucleosome under zero torsion.

**b.** The converted Pol II position from TSS. The shaded region represents an expected nucleosome position. In this assay, it is difficult to resolve the Pol II position inside a nucleosome, so this position is estimated using a linear interpolation of the positions before and after Pol II encounters the nucleosome.

**c.** Mean trajectories of Pol II transcription through a nucleosome. All traces were aligned when Pol II reached the entry of the nucleosome ($t$ = 0), with the shaded regions representing 30% of the standard deviation ($N$ = 234 for -TFIIS; $N$ = 418 for +TFIIS).

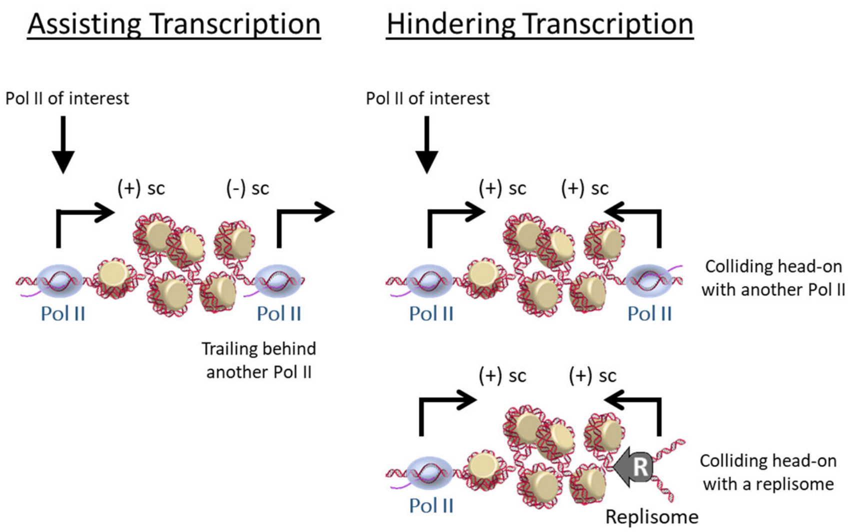


**Supplementary Figure 9.** Cartoon illustrating how processes downstream a transcribing Pol II can modulate the torsion that Pol II encounters. Pol II trailing behind another transcribing Pol II experiences reduced torsional stress. Pol II encountering another Pol II (or a replisome) head-on experiences increased torsional stress.

**Supplementary Table 1**. Primer sequences.

| Oligo Name | Sequence (5' → 3') |
|---|---|
| RNA 14 | UUUUU<u>AUCGAGAGG</u> (Underlined sequence complementary to the gap) |
| ARP77 NTS | TCAGCCCTATAGGATACTTACAGCCATCGAGAGGGACAAGGCGAATA CCCATCCCAATCGGCCTGCTGGTGACACC |
| ARP78 TS | GGTGTCACCAGCAGGCCGATTGGGATGGGTATTCGCCTTGTCCCTCT CGATGGCTGTAAGTATCCTATAGGGCTGA |
| Upstream NTS | GACCTAATTCGAGCTCGGTACCCGGGGATCC |
| Downstream NTS | CAGCGGATCCTCTAGAGTCCTTCAGCGAT |
| Short TS | CTGAAGGACTCTAGAGGATCCGCTGAGGTGTCACCAGCAGGCCGATT GGGATGGGTATTCGCCTTGTCCCTCTCGATGGCTGTAAGTATCCTAT AGGGCTGAGGATCCCCGGGTACCGAGCTCGAATTAG |
| 0.6 kb gapped PCR F | ACTACACCTAGTGCAGGGCGCGTACTATG |
| 0.6 kb gapped PCR R | TAATCCAAATCGATGGCCCGAAGAGTGGGTTTTAC |
| 0.6 kb gapped PCR R Nuc | ATTATACAGGGGCTGACCCGAAGAGTGGGTTTTAC |
| 4.6 kb center segment F | AATACTGTTCACCAACGATCTGGATCACG |
| 4.6 kb center segment R | TTTGAGCGTGGGTCTCGCGGTATCATT |
| 6.5 kb upstream segment F | GCTTCACTCGTGCTTTTGTTCCTTATTTT |
| 6.5 kb upstream segment R | ACGCCAAGCTTCCACATC |
| 0.3 kb NPE segment F | TGCCAAAACAGCCCCTGTATCACTGCG |
| 0.3 kb NPE segment R | AAACCGGAAGACATGTGCTCGTTAGTTGGGTCGAC |
| 12.3 kb center segment F | TTAACTTGGTCTCTGCACGTTTTCAACAGTGATGAGG |
| 12.3 kb center segment R | ACCTTGCCATCGATTTGGGGTTGTAATAGTTTATCCG |

**Supplementary Movie 1.** Real-time visualization of Pol II rotation of DNA.

This video animation shows an example trace of Pol II rotation of DNA under +3.2 pN·nm resistance torque as shown in Fig. 1b. The Pol II rotation of DNA is visualized via the rotation of the trapped nanofabricated quartz cylinder in the AOT, as the cylinder rotates to follow DNA rotation. The inset circle represents the top view of the cylinder with its angular orientation indicated by the red arrow. Only a portion of this trace (from 25 s to 40 s) is animated. As shown, Pol II continuously rotates the DNA (red regions of the curve) and is interrupted by brief pauses (black regions of the curve). The scale bar provides the conversion to distance transcribed.
